# Measurements of maize root plasticity under water stress in hydroponic chambers

**DOI:** 10.1101/380592

**Authors:** Talukder Z. Jubery, Sisi Liu, Thomas Lubberstedt, Baskar Ganapathysubramanian, Daniel Attinger

## Abstract

Under water stress, plants adjust root traits including depth of root system, root diameter, density of root per volume of soil, hydraulic conductance of root. In this experimental study, we present a method to quantify how hydraulic traits of maize roots adapt to drought. The experiments involve a microfluidic flow sensor and a custom-built pressure chamber, made of transparent plastic for visualization purposes. We measured how maize genotypes (PHB47 and PHZ51) grown for a week in deionized (DI) water and one day in hydroponic nutrients solution (called the irrigated condition) respond to one week of water stress. Conditions of water stress (called drought conditions) were created by mixing Polyethylene Glycol with the nutrients solution. Results show that under drought, the roots of both genotypes respond by approximately halving their global hydraulic conductance. This adjustment seems to be achieved mainly by reductions of the total surface area of the roots. Interestingly, the measured hydraulic conductivity of the roots grown under drought was significantly larger. In all, this study sheds light on how plants adapt to water stress in a hydroponic system, by decreasing root area and increasing root permeability.

## 2. Introduction

Climate change and population growth incite both scientists and engineers to investigate methods to increase food production. Climate change increases drought-index (insufficient soil moisture level to meet the plant needs for water as defined in [1]) all over the world, and in the near future we would have to increase yield and decrease water use.

Improved performance of crops under water stress [2], and conservative or efficient water use would have a vast impact on the sustainability of agriculture, especially in the 1.3 billion hectares of marginal agricultural land [3] globally. In arid regions where irrigation water is at a premium, conservative crop water use would allow the cultivation of a broader range of crops, even with just percentage decreases in water use. In developing countries, such as those in sub-Saharan Africa, conservative water use would be beneficial as it would increase the probability of crops surviving typical dry periods during the growing season [4]. Plant’s root system size, properties, and distribution strongly influence it’s access to water, and plants respond to water stress by adjusting root traits rooting density [5], root length[5, 6], root diameter [7], along with other physiological traits [8]. Therefore, it is imperative to find a way to track and quantify root system attributes in assessing desirable root traits that could improve plant performance under drought condition.

Among various important root traits, root depth, root diameter, rooting density, hydraulic conductance, hydraulic conductivity are found to be regulated by plants under drought [9, 10]. Tracking and quantification of root traits including root depth, root diameter, rooting density are performed by 2D/3D imaging of the root system and subsequent image analysis. On the other hand, several methods are used to quantify hydraulic conductance namely, root pressure probe [11], and high-pressure flow meter [12], pressure chamber [13]. Qing-ming et al. showed that [14] there were no significant differences among hydraulic conductance values of maize seedlings measured by these methods.

In their seminal work, J. Passioura et al. [13] developed a pressure chamber encapsulating the root of a plant, while the stem and the shoots were outside of the chamber. Measurements of the pneumatic pressure applied inside the chamber and of transpiration rates and water flow rates measured on cut leaves allow the determination of the hydraulic conductance of the plant. So-called Passioura chambers are typically metallic and do not allow optical access to the root. Here we develop a Passioura chamber with optical access to the roots, and a microfluidic sensor mounted on the stem for measuring water transport. This optical access could facilitate 3D imaging [15] while the root system is in the chamber.

In this manuscript, we report on the design and manufacturing of a transparent pressure chamber, as well as two main results describing how maize adapts to drought under hydroponic conditions.

## 3. Materials and methods

### Plant cultivation and artificial drought condition

Measurements were performed on 15-day-old seedlings for two maize genotypes, PHB47 and PHZ51. PHB47 and PHZ51 are two elite expired PVP (Plant Variety Protection) inbred lines which belong to Iowa Stiff Stalk and Non-Stiff Stalk heterotic pools, respectively [Brenner et al. 2012]. Seeds were collected from cold storage, and 50 similar sized seeds are selected by visual inspection. Seeds were sterilized with Clorox solution (6% sodium hypochlorite) for 15 min in a beaker with a magnetic stir and subsequently washed twice using DI water. Seeds were then placed on brown germination papers (Anchor Paper, St. Paul, MN, USA), twenty seeds per paper, and rolled up vertically. Before placing the seeds, the papers were moisureized with fungicide solution Captan (2.5 g/ L). All seed rolls were placed into 2-L glass beakers containing 1.4 L of autoclaved deionized water. The glass beakers were kept in a growth chamber, where growing conditions were maintained as per the protocols in [16] with a 16 h photoperiod, a temperature cycle of 25 °C/22 °C (day/night) and 65% relative humidity, light intensity was 200 μmol photons m^−2^s^−1^. Every other day the beakers were refilled with water to maintain the water level at 1.4 L.

After seven days, the seedlings were taken out of the roll, and roots with similar length were selected for transplantation. The selected seedlings were transplanted individually in two custom-made hydroponic systems as shown in **Fig 1** with static nutrient solution and continuous aeration, corresponding to water potential of about 0 MPa (our control case). The solution was prepared by mixing eleven chemical compounds (Calcium nitrate (Ca(No_3_)_2_·4H_2_O), potassium sulfate (K_2_SO_4_), Magnesium sulfate (MgSO_4_·7 H_2_O), potassium chloride (KCl), Monopotassium phosphate (KH_2_PO_4_), Boric acid (H_3_BO_3_), Manganese II sulfate (MnSO_4_·H_2_O), Copper sulfate-5 hydrate (CuSO_4_·5H_2_O), Zinc sulfate-7 hydrate (ZnSO_4_·7H_2_O), Ammonium molybdate ((NH_4_)_6_Mo_7_O_24_·4H_2_O), Iron chelate(Fe-EDTA) with water as described in [17].

**Fig 1.**
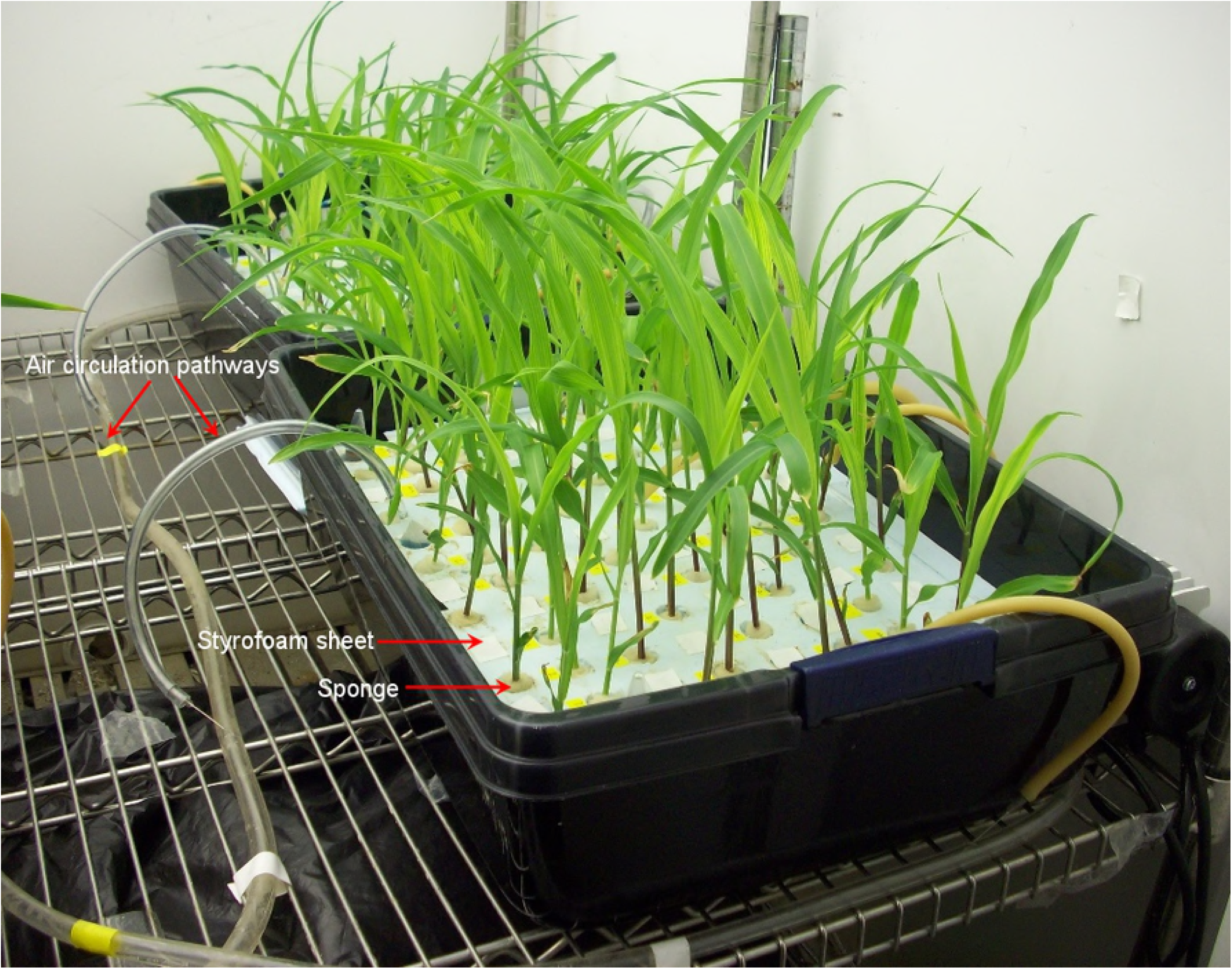
Custom-built hydroponic systems were used to grow transplanted seedlings. The seedlings were individually transplanted via a Styrofoam sheet and pieces of sponge. First through holes were made on the sheet and then the seedlings were transplanted through the holes and held with a piece of sponge. In the system, continuous aeration was introduced using an air pump.

Initially, both hydroponic systems contain the only nutrient solution. After 24 hours, water stress was created in one of the hydroponic systems by adding 15% (w/v) Polyethylene Glycol (PEG) 6000M in the nutrient solution [18], corresponding to water potential of −0.3 MPa. Then, after another seven days, seedlings were removed from the growth chamber for measuring the hydraulic conductance with the pressure chamber described below, and extract root traits with the software ARIA [19].

### Design of pressure chamber and flow sensor

We designed and assembled a pressure chamber to measure the hydraulic conductance of whole root systems (see S1 Appendix for the detail). The device is based on the root transport model described in [13], where the water flow rate J=L(ΔP-σΔΠ), and, L is the root conductance [m^3^/Pas], ΔP=P_1_−P_2_ is the difference of hydrostatic pressure, including gravity, and ΔΠ is a difference of osmotic pressure, with σ is the reflection coefficient. In the device the gradient of water potential (ΔP-σΔΠ) is in the natural direction, that is, the potential is higher around the root system than that at the stem.

### Measurement of hydraulic conductance

For the measurement, a plant was cut at the stem just above the base of the root system and inserted in the pressure chamber as shown in **Fig 2**. The pressure chamber was filled with ultrapure deionized (DI) water (Milipore, US). The differential pressure (ΔP) across the root system and the end of the cut stem was controlled and measured with a pressure controller (Alicat, PC-Series). A transparent tubing was fitted to the cut stem and another end of the tube was connected to a flow sensor. The flow rate, J, through the tube was measured with the flow sensor or calibrated fluidic resistance. The hydraulic conductance (L) was measured by using J = L(ΔP-σΔΠ), where ΔΠ was considered zero, as only DI water was used in our system. A detailed description of the apparatus and of the experimental protocols are included in the supplementary information.

**Fig 2.**
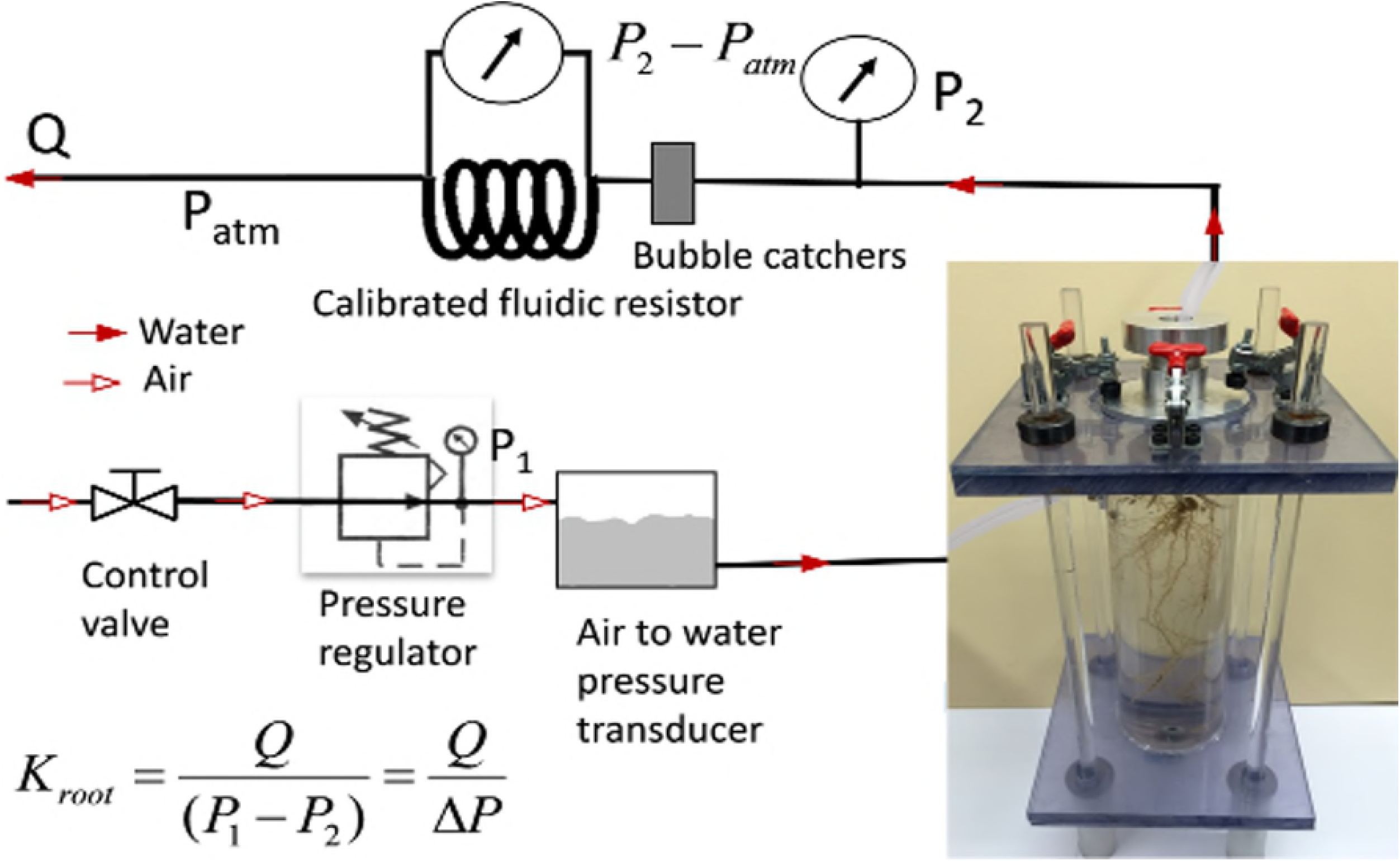
Experimental setup of the flow sensor. The pressure chamber was filled with DI water. A custom-made compression gasket-fitting was used to hold the cut stem and ensure leak proof. The end of the cut stem was connected to a calibrated microfluidic tube (thick black line), the pressure drop across the calibrated tubing was measured via a differential pressure gauge (P_2_-P_atm_). The pressure drop across the root system was measured via another pressure gauge (P_1_-P_2_).

### Measurement of root traits from digital image

After hydraulic conductance measurement, the root systems were imaged using a high-resolution scanner, EPSON Expression 10000 XL scanner system (Copyright 2000-2014 Epson America, Inc). During the imaging the roots were spread out as much as possible to remove overlapping. Using ARIA [20], 27 different visible traits including total surface area, root length, were measured from the image. The surface area was evaluated by considering root system had a circular cross-section where the diameter was the local width of the root segment observed in the image.

## 4. Results

First we validated the accuracy, repeatability, and fidelity of our measurements. We, then, quantified and compared hydraulic conductance of the root system for the genotypes, PHB47 and PHZ51. Finally, we measured the total surface area, total length of the root systems using ARIA and investigated the adaptation of the hydraulic conductivity between the genotypes.

### Validation of the hydraulic conductance measurement

To validate the accuracy, repeatability, and fidelity of our system, we measured the global root conductance of the hydroponically grown seedlings of an arbitrary maize genotype. The seedlings were grown in irrigated condition ((pure water and nutrient solutions). **Fig 3** indicates the global conductance measurement of three samples: two two-week-old seedlings (samples 1 and 2), one three-week-old seedling (sample 3). The measured conductance of two-week-old seedlings are 7.4×10^−15^ [m3/Pa.s] and 6.6 x10^−15^ [m3/Pa.s], respectively, and that of the three-week-old is 5.1 x10^−14^ [m3/Pa.s]. The repeatability and precision of the measurements are very good as shown by the error bars in **Fig 3**. The measured global root conductance of two-week-old seedlings is consistent with the range of published values [21], from 6.6 to 7.7 x10^−15^ [m3/Pa.s].

**Fig 3.**
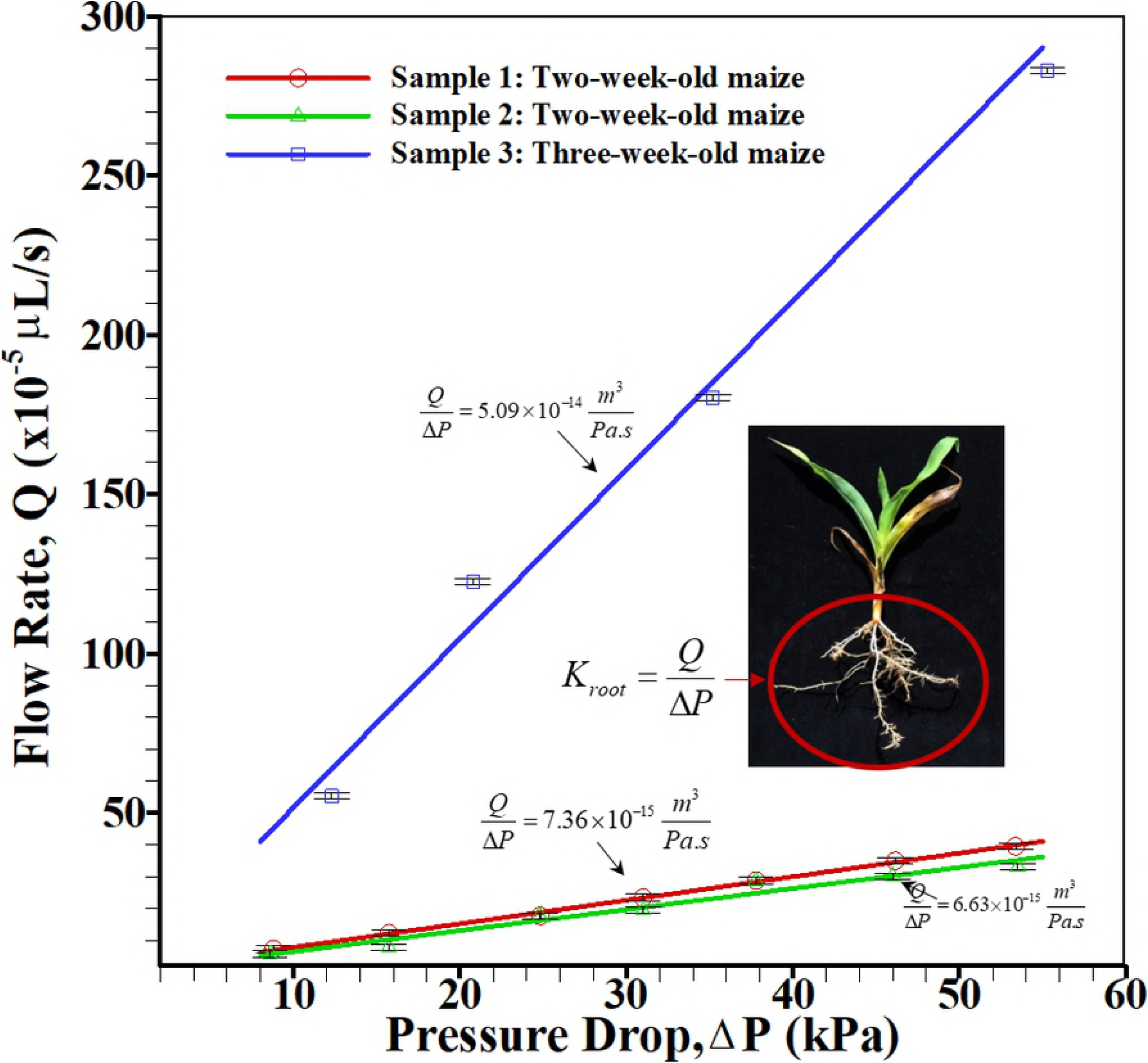
Validation of the flow sensor measurement. Flow rates through the root system of two- and three-week-old seedlings were measured at different applied pressures. Each measurement was repeated ten times. The error bars indicate the standard deviation among the flow rate measurements. The global hydraulic conductance was evaluated from the slope of the best-fitted line passing through the pressure drop vs. mean flow rate measurements. The measured conductance of two-week-old seedlings is consistent with the range of published values [17].

### Adaptation of root traits under drought

Our measurements indicate that seedlings cultivated for seven days under drought (water stress condition) typically exhibit a lower hydraulic conductance than those cultivated in irrigated condition (pure water and nutrient solutions) (**Fig 4** (A)). The reduction of hydraulic conductance of the genotypes PHB47 and PHZ51 are, respectively, 53% and 60%. Reduction in hydraulic conductance indicates plant become water conservative, uses less water during drought. They had similar total root volume (via eyeballing), after the initial first week in the germination paper, but after the second week in different water stress conditions, the total root volumes were different. Under drought, the total root volumes were smaller than those in the irrigated conditions for both genotypes (**Fig 4** (B)), i.e. the root system has less surface area and pathways to uptake water. The overall root surface area under drought was also found around 70% smaller than that in normal/irrigated condition.

**Fig 4.**
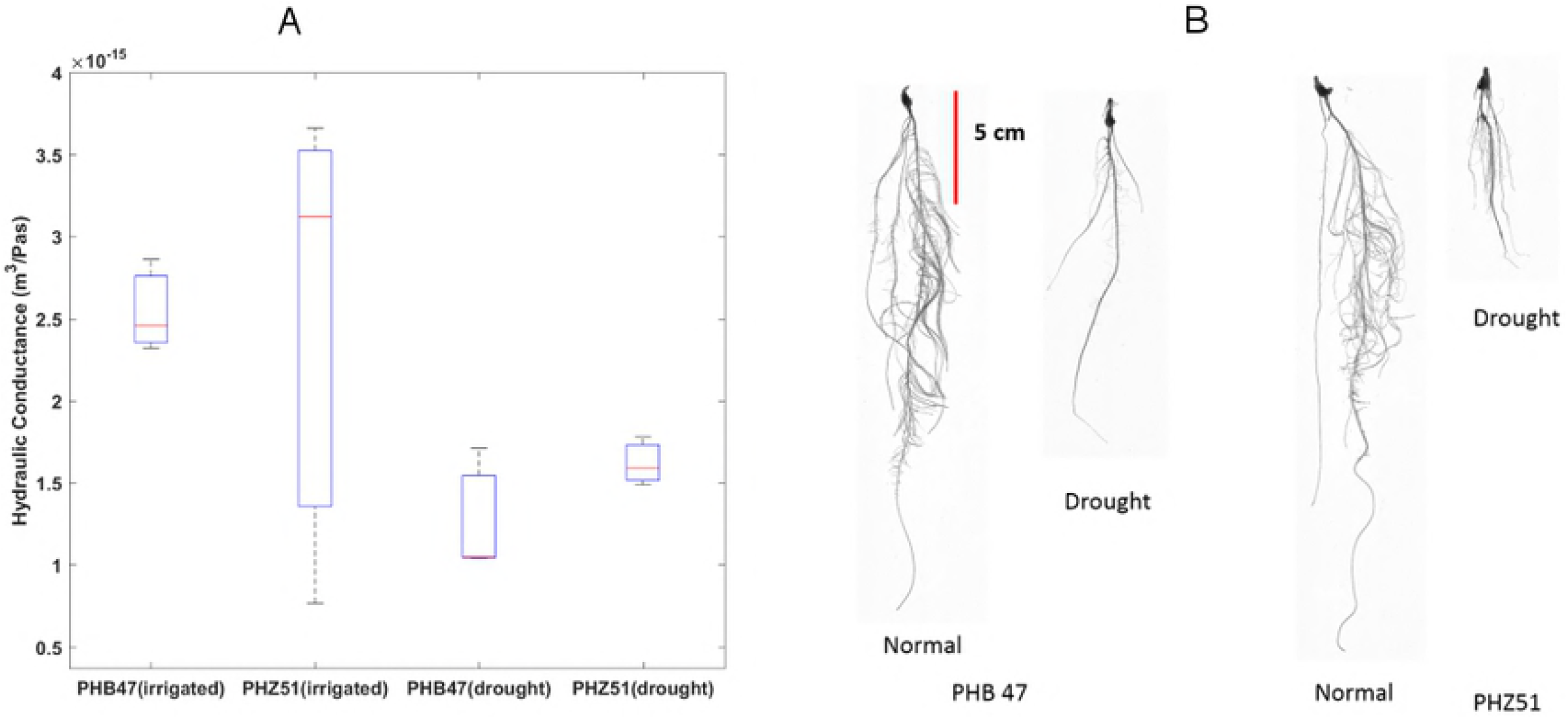
(A) Overall hydraulic conductance of the genotypes under normal/irrigated and drought conditions, the box plots were generated from three data set, (B) photograph of the genotypes in normal/irrigated and drought conditions.

The overall hydraulic conductivity [22] of the genotypes were estimated by dividing overall hydraulic conductance with overall surface area. The hydraulic conductivity under drought was increased significantly for both genotypes (**Fig 5**), that is 50% for PHB47 and 150% for PHZ51. Similar trends also reported for young maize seedlings grown in pots by Zhang et al. [23]. They found an increased hydraulic conductivity when plants grown under drought conditions were re-watered. The adaptation of the hydraulic conductivity could be due to the change in the structural components in roots and/or increased abundance and conductance of aquaporins [24–28].

**Fig 5.**
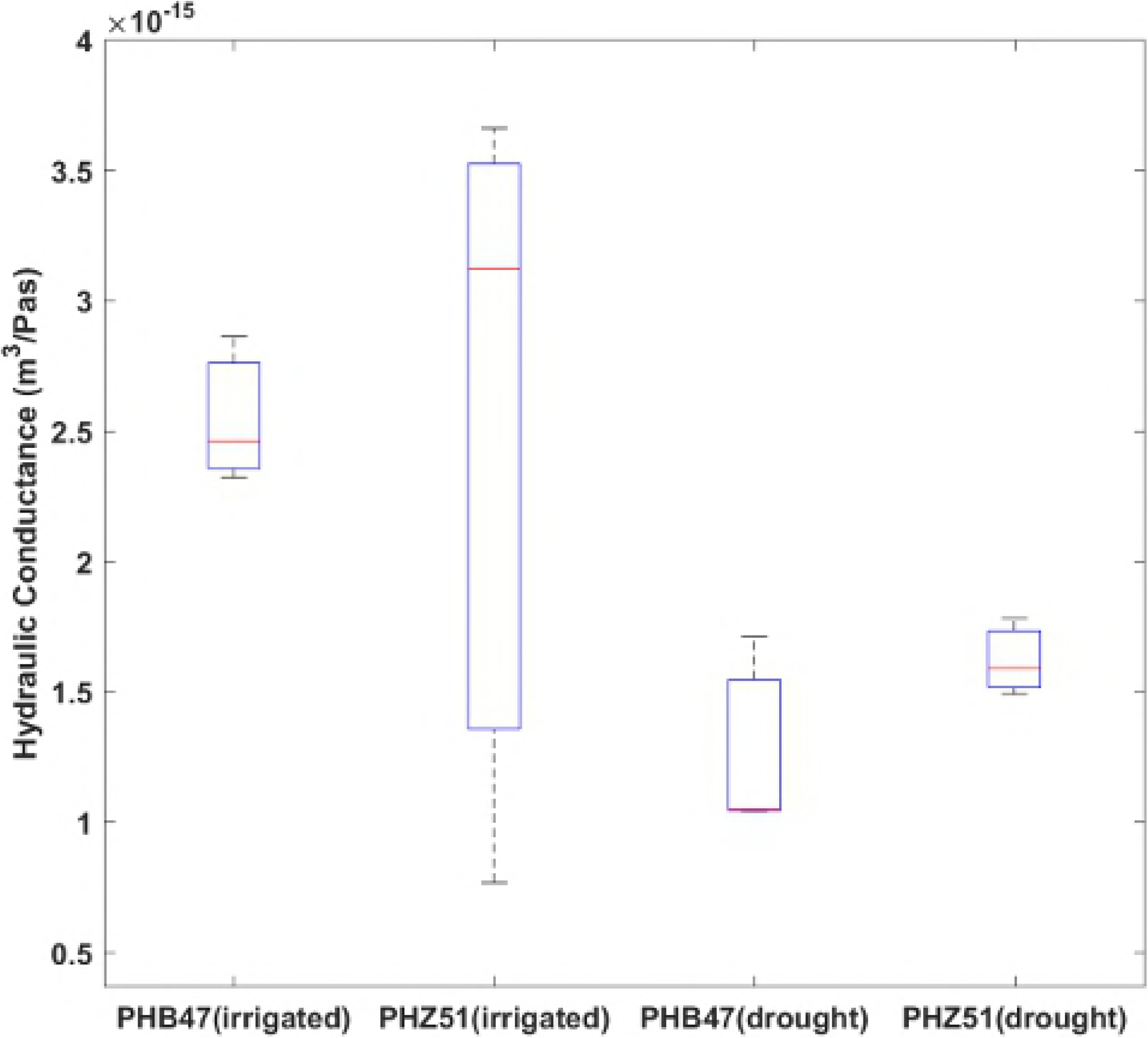
Hydraulic conductivity of plants under normal and drought conditions.

## 5. Conclusions

In this paper, we present a methodology to quantify the adaptation of maize root hydraulic traits to the water stress/drought. The experiments utilize microfluidic flow sensors and a custom-built pressure chamber. The pressure chamber was made of transparent material for visualization purposes. Two inbred maize genotypes (PHB47 and PHZ51) were used to investigate the root systems adaption to artificial drought condition. The drought condition was created by mixing Polyethylene Glycol (PEG) 6000 with the nutrients solution. In response to drought, roots of both genotypes adapted similarly — the roots grew smaller in size compared to the roots in irrigated condition, i.e. the overall surface area of roots was reduced. The surface area was measured by an in-house quantitative phenotyping software. Interestingly, the measured hydraulic conductivity of the roots grown under drought is significantly larger than that of roots grown under irrigated conditions. Similar observations were reported for several genotypes of maize in the responses to daily vapor pressure demand [29] and moderate water stress [30]. In all, this study sheds light on how plants adapt to water stress in a hydroponic system, by decreasing root area and increasing root hydraulic conductivity. The developed methodology can be used to investigate plant responses in other abiotic stresses e.g., salt, heat.

## Acknowledgements

### 6. Acknowledgment

The authors gratefully acknowledge financial support from the Presidential initiative for interdisciplinary research at Iowa State University. The authors thank Matthew Gilbert, UC Davis for his valuable suggestions in building the experimental setup.

## 7. Supporting information

**S1 Appendix. Design and fabrication of pressure chamber.**

**S2 Appendix. Design and calibration of microfluidic pressure sensor.**

**S3 Appendix. Sample loading for the measurement of hydraulic conductance.**

**S1 Fig.**
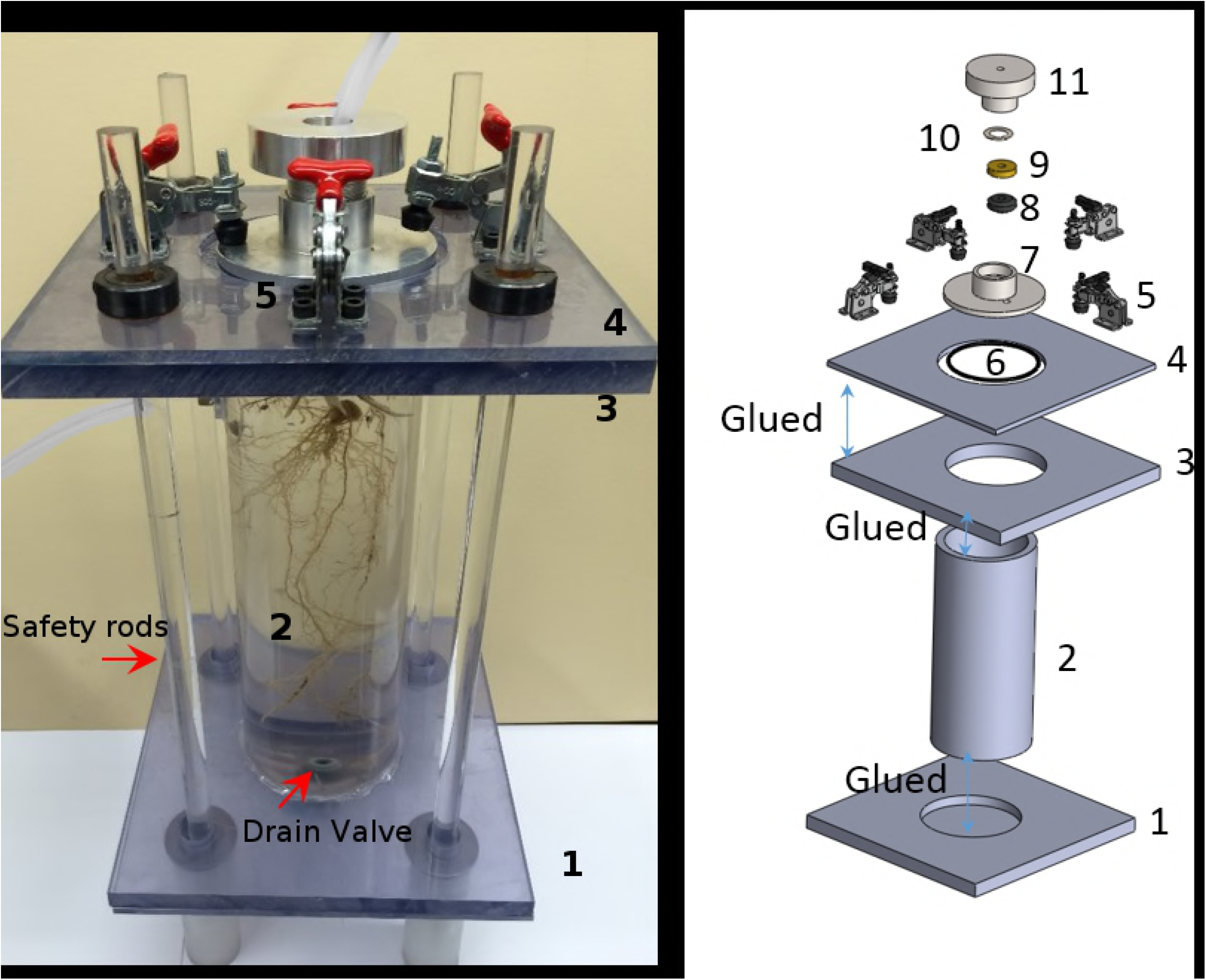
Photograph and simplified exploded view of the pressure chamber.

**S2 Fig.**
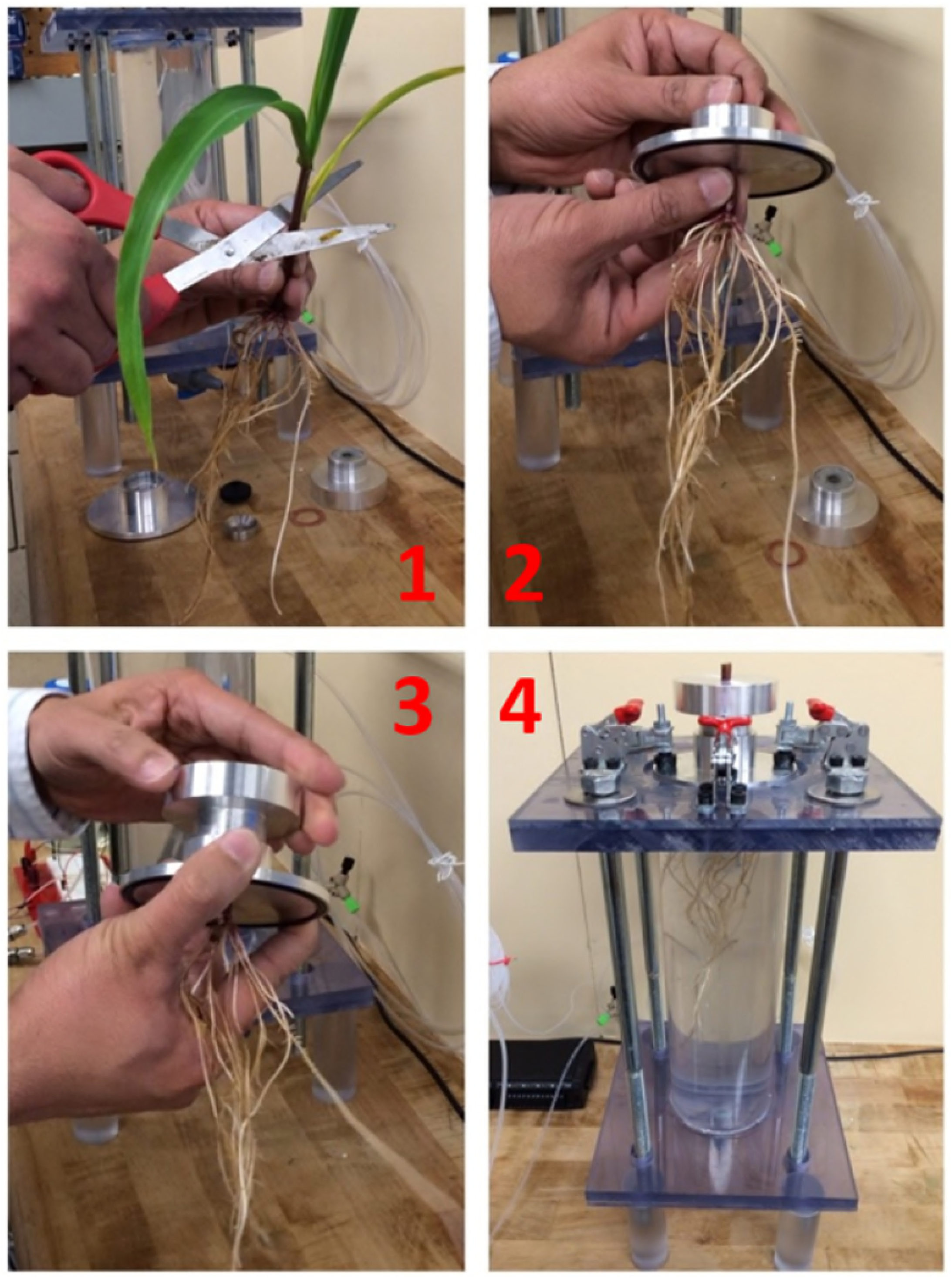
Sequential images of the sample loading procedure in the chamber.

## References

1. Catherine L. Kling, Raymond W. Arritt, Gray Calhoun, David A. Keiser, John M Antle, Jeffery Arnold, et al. Research Needs and Challenges in the FEW System: Coupling economic models with agronomic, hydrologic, and bioenergy models for sustainable food, energy, and water systems: A white paper prepared for the National Science Foundation’s Food, Energy, and Water Workshop held at Iowa State University, October 11–12, 2015. March 2016.

2. Araus J, Slafer G, Reynolds M, Royo C. Plant breeding and drought in C3 cereals: what should we breed for? Annals of botany. 2002;89(7):925–40.

3. Post WM, Nichols JA, Wang D, West TO, Bandaru V, Izaurralde RC. Marginal lands: concept, assessment and management. Journal of Agricultural Science. 2013;5(5):p129.

4. Sinclair TR, Marrou H, Soltani A, Vadez V, Chandolu KC. Soybean production potential in Africa. Global Food Security. 2014;3(1):31–40.

5. Zhan A, Schneider H, Lynch J. Reduced lateral root branching density improves drought tolerance in maize. Plant Physiology. 2015:pp. 00187.2015.

6. Lu ZJ, Neumann PM. Water stress inhibits hydraulic conductance and leaf growth in rice seedlings but not the transport of water via mercury-sensitive water channels in the root. Plant Physiology. 1999;120(1):143–51. doi: 10.1104/pp.120.1.143. PubMed PMID: WOS:000080329200016.

7. Comas LH, Becker SR, Cruz VMV, Byrne PF, Dierig DA. Root traits contributing to plant productivity under drought. Frontiers in Plant Science. 2013;4. doi: 10.3389/fpls.2013.00442. PubMed PMID: WOS:000331467000001.

8. McNear Jr D. The rhizosphere-roots, soil and everything in between. Nature Education Knowledge. 2013;4(3):1.

9. Hammer GL, Dong Z, McLean G, Doherty A, Messina C, Schussler J, et al. Can changes in canopy and/or root system architecture explain historical maize yield trends in the US corn belt? Crop Science. 2009;49(1):299–312.

10. Khonghintaisong J, Songsri P, Toomsan B, Jongrungklang N. Rooting and Physiological Trait Responses to Early Drought Stress of Sugarcane Cultivars. Sugar Tech. 2018;20(4):396–406. doi: 10.1007/s12355-017-0564-0. PubMed PMID: WOS:000431412400004.

11. Steudle E, Oren R, Schulze E-D. Water transport in maize roots measurement of hydraulic conductivity, solute permeability, and of reflection coefficients of excised roots using the root pressure probe. Plant Physiology. 1987;84(4):1220–32.

12. Tyree MT, Patiño S, Bennink J, Alexander J. Dynamic measurements of roots hydraulic conductance using a high-pressure flowmeter in the laboratory and field. Journal of experimental botany. 1995;46(1):83–94.

13. Passioura JB, Munns R. Hydraulic resistance of plants. II. effects of rooting medium, and time of day, in barley and lupin. Australian Journal of Plant Physiology. 1984;11(5):341–50. PubMed PMID: WOS:A1984TW88600002.

14. Li Q-m, Liu B-b. Comparison of Three Methods for Determination of Root Hydraulic Conductivity of Maize (Zea mays L.) Root System. Agricultural Sciences in China. 2010;9(10):1438–47. doi: 10.1016/s1671-2927(09)60235-2. PubMed PMID: WOS:000283106400006.

15. Clark RT, MacCurdy RB, Jung JK, Shaff JE, McCouch SR, Aneshansley DJ, et al. Three-Dimensional Root Phenotyping with a Novel Imaging and Software Platform. Plant Physiology. 2011;156(2):455–65. doi: 10.1104/pp.110.169102. PubMed PMID: WOS:000291146800002.

16. Abdel-Ghani AH, Kumar B, Reyes-Matamoros J, Gonzalez-Portilla PJ, Jansen C, San Martin JP, et al. Genotypic variation and relationships between seedling and adult plant traits in maize (Zea mays L.) inbred lines grown under contrasting nitrogen levels. Euphytica. 2013;189(1):123–33. doi: 10.1007/s10681-012-0759-0. PubMed PMID: WOS:000312127200009.

17. Hoagland DR, Arnon DI. The water-culture method for growing plants without soil. Circular California Agricultural Experiment Station. 1950;347(2nd edit).

18. Lian H-L, Yu X, Lane D, Sun W-N, Tang Z-C, Su W-A. Upland rice and lowland rice exhibited different PIP expression under water deficit and ABA treatment. Cell research. 2006;16(7):651–60.

19. Brenner EA, Blanco M, Gardner C, Lübberstedt T. Genotypic and phenotypic characterization of isogenic doubled haploid exotic introgression lines in maize. Molecular breeding. 2012;30(2):1001–16.

20. Pace J, Lee N, Naik HS, Ganapathysubramanian B, Lübberstedt T. Analysis of Maize (Zea mays L.) Seedling Roots with the High-Throughput Image Analysis Tool ARIA (Automatic Root Image Analysis). PLoS One. 2014;9(9):e108255. doi: 10.1371/journal.pone.0108255.

21. Mu Z, Zhang S, Zhang L, Liang A, Liang Z. Hydraulic conductivity of whole root system is better than hydraulic conductivity of single root in correlation with the leaf water status of maize. Botanical Studies. 2006;47(2):145–51.

22. North G, Nobel P. Heterogeneity in water availability alters cellular development and hydraulic conductivity along roots of a desert succulent. Annals of Botany. 2000;85(2):247–55.

23. Zhang JH, Zhang XP, Liang JS. Exudation rate and hydraulic conductivity of maize roots are enhanced by soil drying and abscisic-acid treatment. New Phytologist. 1995;131(3):329–36. doi: 10.1111/j.1469-8137.1995.tb03068.x. PubMed PMID: WOS:A1995TH78300005.

24. Holbrook NM, Zwieniecki MA. Vascular transport in plants: Academic Press; 2011.

25. Parent B, Hachez C, Redondo E, Simonneau T, Chaumont F, Tardieu F. Drought and Abscisic Acid Effects on Aquaporin Content Translate into Changes in Hydraulic Conductivity and Leaf Growth Rate: A Trans-Scale Approach. Plant Physiology. 2009;149(4):2000–12. doi: 10.1104/pp.108.130682. PubMed PMID: WOS:000264796900035.

26. Vandeleur RK, Mayo G, Shelden MC, Gilliham M, Kaiser BN, Tyerman SD. The Role of Plasma Membrane Intrinsic Protein Aquaporins in Water Transport through Roots: Diurnal and Drought Stress Responses Reveal Different Strategies between Isohydric and Anisohydric Cultivars of Grapevine. Plant Physiology. 2009;149(1):445–60. doi: 10.1104/pp.108.128645. PubMed PMID: WOS:000262261500047.

27. Kaldenhoff R, Grote K, Zhu JJ, Zimmermann U. Significance of plasmalemma aquaporins for water-transport in Arabidopsis thaliana. Plant Journal. 1998;14(1):121–8. doi: 10.1046/j.1365-313X.1998.00111.x. PubMed PMID: WOS:000073317600013.

28. Laur J, Hacke UG. Transpirational demand affects aquaporin expression in poplar roots. Journal of Experimental Botany. 2013;64(8):2283–93. doi: 10.1093/jxb/ert096. PubMed PMID: WOS:000319433200014.

29. Caldeira CF, Bosio M, Parent B, Jeanguenin L, Chaumont F, Tardieu F. A Hydraulic Model Is Compatible with Rapid Changes in Leaf Elongation under Fluctuating Evaporative Demand and Soil Water Status. Plant Physiology. 2014;164(4):1718–30. doi: 10.1104/pp.113.228379. PubMed PMID: WOS:000334342800016.

30. Yang T, Liang Z, Xue J, Kang S. Diversity of water use efficiency in various maize varieties. Transactions of the Chinese Society of Agricultural Engineering. 2005;21(10):21–5. PubMed PMID: CABI:20063024619.

